# The isoniazid analog IQG-607 is not a direct substrate for the *Mycobacterium tuberculosis* catalase-peroxidase KatG

**DOI:** 10.1101/2024.09.05.611129

**Authors:** Laísa Quadros Barsé, Candida Deves Roth, Adilio da Silva Dadda, Raoní Scheibler Rambo, Pedro Ferrari Dalberto, Kenia Pissinate, José Eduardo Sacconi Nunes, Renata Jardim Etchart, Pablo Machado, Luiz Augusto Basso, Cristiano Valim Bizarro

## Abstract

Tuberculosis (TB) is an infectious disease caused mainly by *Mycobacterium tuberculosis* (Mtb) and is responsible for millions of deaths. New Mtb strains resistant to TB drugs are emerging and spreading. The first-line TB drug, isoniazid (INH), must be activated inside mycobacterial cells by the catalase-peroxidase enzyme KatG to exert its antimicrobial activity, and mutations on the *katG* gene are a significant cause of INH resistance in clinics. The metal-containing compound IQG-607 is an INH analog developed to inhibit the target of INH, the FASII enzyme enoyl-ACP-reductase (InhA), without requiring KatG. However, we recently showed that inside mycobacterial cells, IQG-607 activity depends on KatG. Hence, this compound might also be activated by KatG to inhibit InhA. We evaluated whether recombinant MtKatG uses IQG-607 as a substrate in oxidation reactions and adduct formation with NAD^+^. A recombinant MtKatG was produced in *E. coli* and purified in a 3-step protocol to obtain a homogeneous protein. An HPLC method was optimized to monitor both oxidation and adduct products, and our assay system was validated by performing control reactions using INH as a substrate. We found that the metal-based compound IQG-607 is not a substrate for recombinant MtKatG under all conditions tested.

## Introduction

Tuberculosis (TB) is a well-known infectious disease, mainly caused by *Mycobacterium tuberculosis* (Mtb), an aerobic pathogenic bacterium that primarily infects the lungs [1]. TB was responsible for millions of deaths when there were no adequate treatments for patients. Even today, TB is one of the 10 leading causes of death globally. In 2022, according to the World Health Organization (WHO), 1.3 million people died from TB [2]. Moreover, the COVID-19 pandemic impacted both diagnosis and treatment of TB [3]. Currently, standard TB treatment consists of an intensive initial phase lasting two months and a continuous phase for an additional four months. The suggested treatment for new TB patients consists of a two-month phase using isoniazid (INH), rifampicin (RIF), pyrazinamide (PZA), and ethambutol (EMB), followed by a four-month phase with INH and RIF [2]. Alternatively, the continuation phase may extend to six months of INH and EMB [2]. TB is a curable disease, and most patients that follow the treatment are cured. However, the emergence and spread of Mtb strains resistant to first-line drugs represent one of the main challenges in treating TB today. Only three new drugs were approved for TB treatment in the last decade: bedaquiline (BDQ), delamanid (DLM) and pretomanid (PMD) [4].

Long treatment periods coupled with the adverse effects of first-line drugs result in high treatment dropout rates, which contribute to the selection and dissemination of resistant strains [5]. Mtb strains resistant to both INH and RIF are defined as multidrug-resistant (MDR strains). Patients affected by TB caused by MDR strains (MDR-TB) need alternative therapies using second-line drugs. Examples are fluoroquinolones and injectable drugs such as amikacin, kanamycin, and capreomycin. However, second-line drugs are more expensive, less effective, and more toxic. In addition, a more extended treatment period (at least 18 months) is also required. The occurrence of extensively drug-resistant (XDR-TB) strains, which are MDR-TB strains that are also resistant to fluoroquinolones and at least one of the second-line injectable drugs, makes the situation even worse, both for TB treatment and eradication [6].

INH resistance alone or in combination with other drugs is the second most common type of resistance observed among TB patients, with rifampicin resistance alone (RR-TB) being the first one. As the most used drug in the clinic, INH resistance represents one of the main current challenges in treating TB [7]. INH has a simple structure consisting of a hydrazide group bounded to a pyridine ring, these two components being essential for its activity (Fig. 1a). High bactericidal activity, low cost, high bioavailability, excellent intracellular penetration, and a narrow spectrum of action make INH one of the most effective antimicrobial agents to treat TB [8]. The most accepted mechanism of action for INH, described in the 1990s, posits that INH acts as a prodrug, requiring intracellular activation of a catalase-peroxidase from Mtb (MtKatG) to exert its activity. When activated, INH is converted to an isonicotinoyl radical, which reacts with nicotinamide adenine dinucleotide (NAD^+^), leading to the INH-NAD adduct [9]. This adduct turns into an active compound that inhibits the NADH-dependent trans-2-enoyl-ACP reductase enzyme (InhA). InhA is an essential enzyme involved in the mycobacterial cell wall and fatty acid biosynthesis. Its inhibition blocks fatty acid elongation, leading to the accumulation of long-chain fatty acids and cell death [10]. Some of the most frequent mutations found in clinical isolates associated with INH resistance are in the gene that encodes KatG. One in particular (S315T) is found in more than 50% of Mtb resistant populations [11–14].

**Fig. 1.**
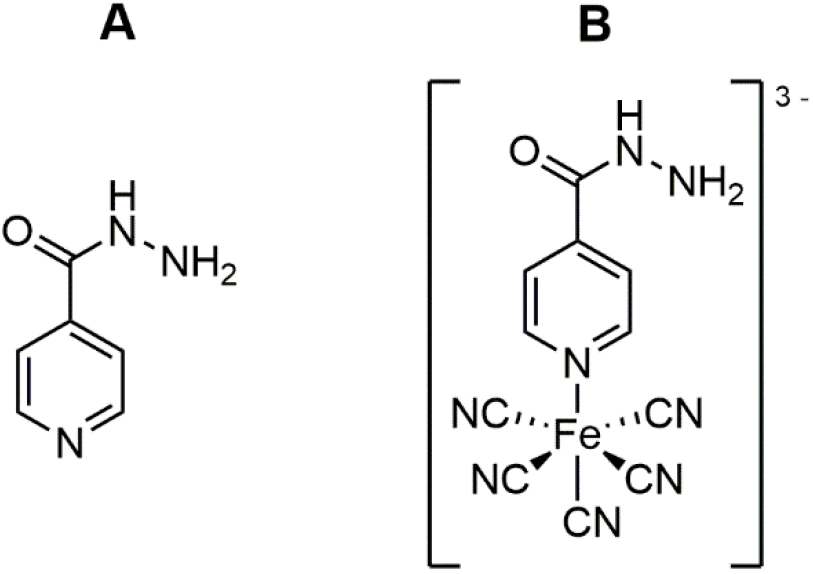
Structure of isoniazid (a) and IQG-607 (b).

There is an urgent need to develop a new drug with anti-tubercular action mainly against resistant strains, with low toxicity, and able to shorten treatment of latent TB infection [15]. Metal-based drugs approved for clinical use (cisplatin, nitroprusside, silver sulfadiazine, and others) [16] share some advantages when compared to organic compounds, such as adjustable redox potential and the ability to build new structures due to a larger number of stereoisomers [17–19]. IQG-607 is a metal-based compound containing a pentacyanoferrate(II) attached to the nitrogen atom of the heterocyclic ring of INH, having the molecular formula [Fe(II)(CN)_5_(INH)]^3-^ (Fig. 1b). This molecule was rationally developed to undergo autoactivation in vivo, eliminating the need for MtKatG activity, as required for INH prodrug activation. In this proposal, the metal center containing the pentacyanoferrate portion would promote electron transfer and form the isonicotinoyl radical, mimicking the activation mechanism of MtKatG [20,21]. This intramolecular autoactivation of IQG-607 would be triggered by reactive oxygen species (ROS), such as hydrogen peroxide, which would oxidize Fe(II) to Fe(III) (I in Fig. 2), and this, in turn, would promote intramolecular oxidation of the isoniazid moiety to isonicotinoyl radical, with Fe(III) being reduced back to Fe(II) (II in Fig. 2). Conceivably, the non-enzymatic formation of the isonicotinoyl radical from IQG-607 would circumvent the need for MtKatG activity inside Mtb cells, as required for INH activation. In this way, this compound could be used as an effective INH surrogate to treat TB cases caused by INH-resistant strains whose mechanism of resistance lies on mutations on the *katG* gene.

**Fig. 2.**
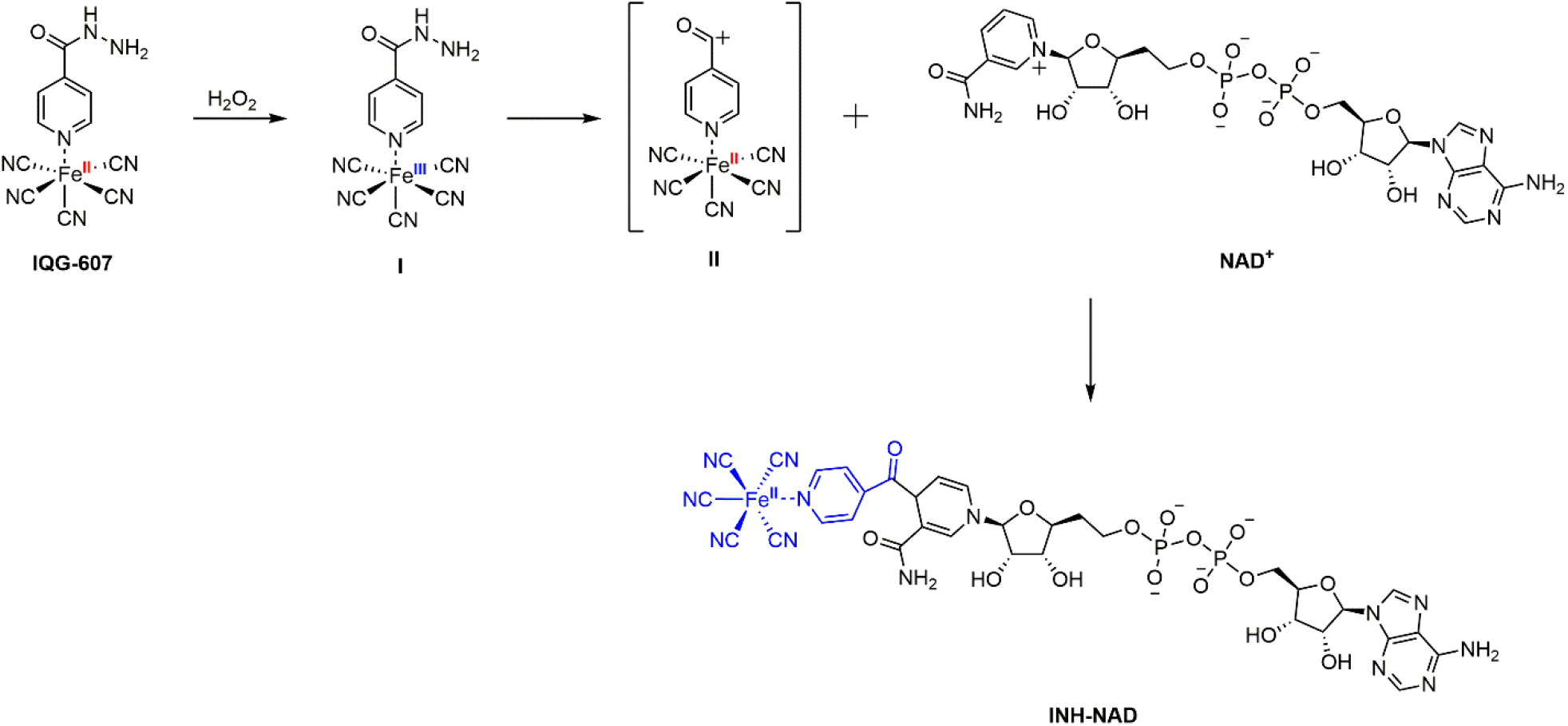
The proposed mechanism through a non-enzymatic oxidative route of IQG-607 in the formation of the isonicotinoyl radical and formation of the complex which inhibits InhA.

Some studies have shown that IQG-607 inhibits in vitro both the InhA WT enzyme and some structural mutants (S94A, I21V, and I47T), known to confer INH’s resistance [15,21-23]. Moreover, lipid extraction and radiolabelling analysis of mycolic acids were performed to study their mechanism of action. Total inhibition of mycolic acid synthesis was observed when IQG-607 was used, like what was found when Mtb cells were treated with INH [22] 25. This result confirmed that the target consisted of the InhA enzyme, corroborating the previously performed in vitro enzyme assays [21,23]. Moreover, we observed a favorable IQG-607 profile in cell culture cytotoxicity without genotoxicity, even at high concentrations, did not induce DNA damage in HepG2 cells and demonstrated an advantage over INH in rats, mice, and mini pigs [15, 24-26].

Recent work from our research group evaluated the mechanism of action of IQG-607 in INH-resistant Mtb strains [27]. Initially, 9 MDR-TB clinical isolates were treated with IQG-607. Unexpectedly, the isolates carrying the *katG*(S315T) mutation (8 out of 9 treated) were found to be resistant to this compound, with MICs ranging from 25 to >100 µg mL^-1^ [27]. Moreover, six spontaneous mutants resistant to the IQG-607 were isolated and found to have alterations in the *katG* gene, four of them with a mutation profile generally unexpected in clinical isolates but common in laboratory-generated mutants [28,29]. Additionally, we established a causal relationship between compound resistance and identified mutations. A knockout strain for the *katG* gene was produced and subsequently complemented with the wild-type (WT) *katG* gene or a variant of that gene encoding the mutant KatG (S315T) (the mutation most often found in INH-resistant clinical isolates). Bacteria complemented with the *katG* (S315T) gene increased its MIC by 64-fold [27]. These results strongly suggest that, inside mycobacterial cells, the antimicrobial action of compound IQG-607 depends on the activity of the MtKatG enzyme, in disagreement with the autoactivation model.

Based on these results, we hypothesized that similarly to INH, IQG-607 needs to be activated by the MtKatG enzyme in the intracellular environment to exert its antimicrobial activity. We evaluated whether IQG-607 is a substrate for MtKatG under different experimental conditions. For this purpose, we produced and biochemically characterized a recombinant MtKatG in *E. coli*. Moreover, we developed analytical methods to detect species formed from INH and IQG-607 compounds under different reaction conditions. We performed enzymatic assays in the absence (air-only background control) or the presence of peroxides as exogenous oxidants (t-BuOOH and H_2_O_2_).

## Materials and methods

### Cloning, expression, purification, oligomeric state, and mass spectrometry identification of recombinant MtKatG

The *M. tuberculosis* catalase-peroxidase (MtKatG) coding gene katG was synthesized by FastBio Ltda and was cloned into the pET23a(+) expression vector using the NdeI and HindIII restriction enzymes. *E. coli* BL21 (DE3)pLysS cells were used as host cells. For more information, see Supplementary Material.

A 3-step purification protocol was developed to obtain homogenous recombinant MtKatG. FPLC was performed using the AKTA system (GE Healthcare), and all purification steps were carried out at 4 °C and sample elution monitored by UV detection at 215, 254, and 280 nm simultaneously. For more information, see Supplementary Material.

For oligomeric state determination, analytical gel filtration was performed using a Superdex 200 HR 10/30 (GE Healthcare) column pre-equilibrated with 50 mM potassium phosphate pH 7.2 containing 150 mM NaCl at a flow rate of 0.6 mL min^−1^, with UV detection at 215, 254, and 280 nm. The LMW and HMW Gel Filtration Calibration Kits (GE Healthcare) were used to prepare a calibration curve. The elution volumes (V_e_) of standard proteins (ferritin, aldolase, coalbumin, ovalbumin, carbonic anhydrase, ribonuclease A, and aprotinin) were used to calculate their corresponding partition coefficient (K_av_, Eq. (1)). Blue dextran 2000 (GE Healthcare) was used to determine the void volume (V_o_). V_t_ is the total bead volume of the column.

The K_av_ value for each protein was plotted against their corresponding logarithm of molecular mass.

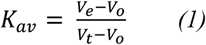

For mass spectrometry identification, MtKatG recombinant protein fractions were analyzed by SDS-PAGE 12%, and a gel slice containing the protein was excised and submitted to in-gel digestion [31,32]. The tryptic digest was separated on an in-house made 20 cm reverse-phase column (5 µm ODSAQ C18, Yamamura Chemical Lab, Japan) using a nanoUPLC (nanoLC ultra 1D plus, Ekisgent, USA), and the eluted peptides transferred to a nanospray ion source connected to a hybrid mass spectrometer (LTQ-XL and LTQ orbitrap Discovery, Thermo, USA). Spectra were searched against a non-redundant *E. coli* BL21(DE3) database and MtKatG sequence with the software COMET [31] in the platform PatternLab for proteomics, and the validity of the peptide-spectra matches were assessed using Paternlab’s module SEPro [33] with a false discovery rate of 1% based on the number of decoys.

### Catalase and peroxidase activity assays

All the measurements were performed using a continuous spectrophotometric assay in quartz cuvettes using a UV-visible Shimadzu spectrophotometer UV2550 equipped with a temperature-controlled cuvette holder. Catalase assay was carried out at 37 °C in 100 mM potassium phosphate pH 7.5. The catalase activity was measured spectrophotometrically by following the consumption of hydrogen peroxide over 60 s at 240 nm (ε_240_ = 0.0435 mM^-1^ cm^-1^) [34], with initial concentrations of H_2_O_2_ ranging from 0.1 to 5 mM, and with a fixed concentration of MtKatG of 8.8 nM. One unit of catalase was defined as the decomposition amount of 1 µmol H_2_O_2_ min^-1^ reaction. Peroxidase assay was carried out at 37 °C in 100 mM potassium phosphate buffer pH 5. The peroxidase activity was measured by following the increase in absorbance for 1 mM 2,2’-azino-bis(3-ethylbenzothiazoline-6-sulfonic acid (ABTS) (ε_406_ = 36.8 mM^-1^ cm^-1^) [35] in the presence of tert-butyl hydroperoxide (t-BuOOH) ranging from 0.25 to 25 mM over 180 s at a fixed enzyme concentration of 25 nM. One unit of peroxidase activity was defined as the oxidation of 1 µmol ABTS min^-1^ reaction. Specific activity was expressed in U/mg of total protein. All assays were performed at least in duplicate. Kinetic parameters (K_m_, V_max_) were obtained from nonlinear regression of Michaelis-Menten plots using Sigma Plot version 12.0.

### INH and IQG-607 oxidation and enzymatic adduct formation

INH and IQG-607 oxidation and adduct formation assays were performed in triplicate at 37 °C as described in previous studies with minor modifications [14,36-38]. Briefly, the reaction sample (1 mL total volume) contained 100 mM potassium phosphate pH 7.5, 1.5 µM MtKatG, 1 mM INH (or IQG-607), 120 µM NAD^+^ and the following oxidants (air only, 400 µM t-BuOOH or 400 µM H_2_O_2_) were used. The total time of reaction was 100 minutes. HPLC analyses were performed using 90 µL injection volume on a Dionex Ultimate 3000 equipped with a diode array detector (DAD) using a reversed-phase C18ec column (Macherey-Nagel, 5 µm particle size, 250 mm x 4.6 mm, Nucleosil) applying a nonlinear gradient from 0 to 15% acetonitrile in 70 mM ammonium acetate solution over 30 min using flowrate at 1 mL min^-1^ and detection was in 260 nm. INH and IQG-607 products of oxidation and adducts were identified by their characteristic UV-visible absorption spectrum as described in the literature. Also, the HPLC fractions containing the four INH-NAD adducts were collected, pooled, and subjected to mass spectrometry analysis using a positive ion mode.

## Results and discussion

### Cloning, expression, purification, oligomeric state, and mass spectrometry identification of recombinant MtKatG

We produced the recombinant MtKatG in a soluble form with an apparent molecular mass of ∼80 kDa. This result agrees with the expected molecular mass of MtKatG (80604.87 Da). A 3-step purification protocol (anionic exchange followed by hydrophobic interaction and lastly size exclusion) was developed and yielded around 5 mg of homogeneous recombinant MtKatG. MtKatG’s (Uniprot code: P9WIE5) identity was confirmed by mass spectrometry, with the identification of 143 unique peptides, 2087 spectral counts, and a sequence coverage of 91% (Fig. S1). By gel filtration chromatography, the recombinant MtKatG was found to have a molecular mass of 176.6 kDa (Fig. S2), indicating that this protein is a homodimer in solution, as previously found [39].

### Catalase and peroxidase activities of recombinant KatG

Before performing INH and IQG-607 oxidation studies, we evaluated the catalase and peroxidase enzymatic activities of recombinant MtKatG. The kinetic parameters (k_cat_, K_m_, and catalytic efficiency k_cat_/K_m_) for the catalase and peroxidase activities are presented in Table 1. The recombinant MtKatG exhibited saturable catalase activity when varying H_2_O_2_ under the tested conditions, yielding a K_m_ of 0.14±0.03 mM and a k_cat_ of 4600±200 s^-1^. The K_m_ reported here is slightly lower than reported in other studies, ranging from 0.6 mM to 30 mM, and the k_cat_ is in agreement with values already reported, ranging from 2800 to 10 000 s^-1^ [11,30,40-43]. Also, the peroxidase activity for one-electron oxidation using a fixed concentration of ABTS and varying t-BuOOH as substrate was performed, and values for K_m_ of 19±6 mM and k_cat_ of 420±70 s^-1^ were obtained. The kinetic constants reported here for peroxidase assay are “apparent” values since the enzyme saturation was not reached under these conditions.

**Table 1.** Kinetic parameters for catalase and peroxidase activities of recombinant MtKatG.

| | $k_{\text{cat}}$ ($\text{s}^{-1}$) | $K_m$ (mM) | $k_{\text{cat}}/K_m$ ($\text{M}^{-1} \text{s}^{-1}$) |
| --- | --- | --- | --- |
| Catalase | 4600±200 | 0.14±0.03 | 33±7 x 10<sup>6</sup> |
| Peroxidase | 420±70 | 19±6 | 22±12 x 10<sup>3</sup> |

### INH and IQG-607 oxidation using MtKatG

As previously described, INH is a substrate of MtKatG, leading to isonicotinic acid (ISA) as a major product and isonicotinamide (INA) as a minor by-product [9]. These oxidized products have no antimicrobial activity against Mtb strains [44]. Using INH as a positive control, we showed that the recombinant MtKatG has a catalytic activity to oxidize INH as expected, forming both ISA and INA (Fig. S3a and Table 2). As a catalase enzyme, MtKatG converts hydrogen peroxide into water and oxygen. As a peroxidase enzyme, it also promotes the oxidation of other compounds, such as INH, through two consecutive oxidation steps [11,30,45]. When peroxides (H_2_O_2_ or t-BuOOH) were added to the oxidation assay, the conversion of INH into ISA and INA was stimulated (Fig. S3b and c and Table 2). Peroxides were used to mimic the macrophage intracellular microenvironment experienced by infecting bacilli since ROS are produced by both the aerobic metabolism of actively replicating Mtb cells and by host cells, which release large amounts of H_2_O_2_ to combat the infection [46]. In the absence of MtKatG in the reaction, no oxidation of INH occurs, even with the addition of peroxides (Fig. S3d), demonstrating that the formation of oxidation products from INH requires its enzymatic activity.

**Table 2.** Percentage of degradation of INH in absence or presence of recombinant MtKatG.

|  | - MtKatG | + MtKatG |
| --- | --- | --- |
| Air | ND* | 42.3±0.2% |
| H<sub>2</sub>O<sub>2</sub> | ND* | 56±1% |
| t-BuOOH | ND* | 56±4% |

\*ND: no degradation.

Differently from the reactions mentioned above using INH, we observed oxidation products when this compound was replaced by IQG-607 (Fig. 3). IQG-607 alone or with peroxides (H_2_O_2_ or t-BuOOH) decomposed to ISA mainly, but not to INH. Based on a previous study, the coordinated pentacyanoferrate(II) favors a pathway for isonicotinic acyl radical formation, generating ISA [47]. Also, IQG-607 demonstrates a band-shift in UV-spectra from ∼445 to ∼420 nm after 100 minutes of reaction. It has already been reported IQG-607 itself presents a well-defined band in 436 nm and its degradation products in 415 nm [48] (Fig. 4). Moreover, the formation of oxidation products observed in our control reactions without the addition of MtKatG (Fig. 3a, c, e, and Table 3) was partly suppressed by the addition of this enzyme (Fig. 3b, d, f, and Table 3). This effect does not depend on different oxidizing agents used (Table 3). So, presumably, MtKatG acts as a reactive oxygen species (ROS) scavenger, removing oxidative agents that promote IQG-607 decomposition.

**Fig. 3.**
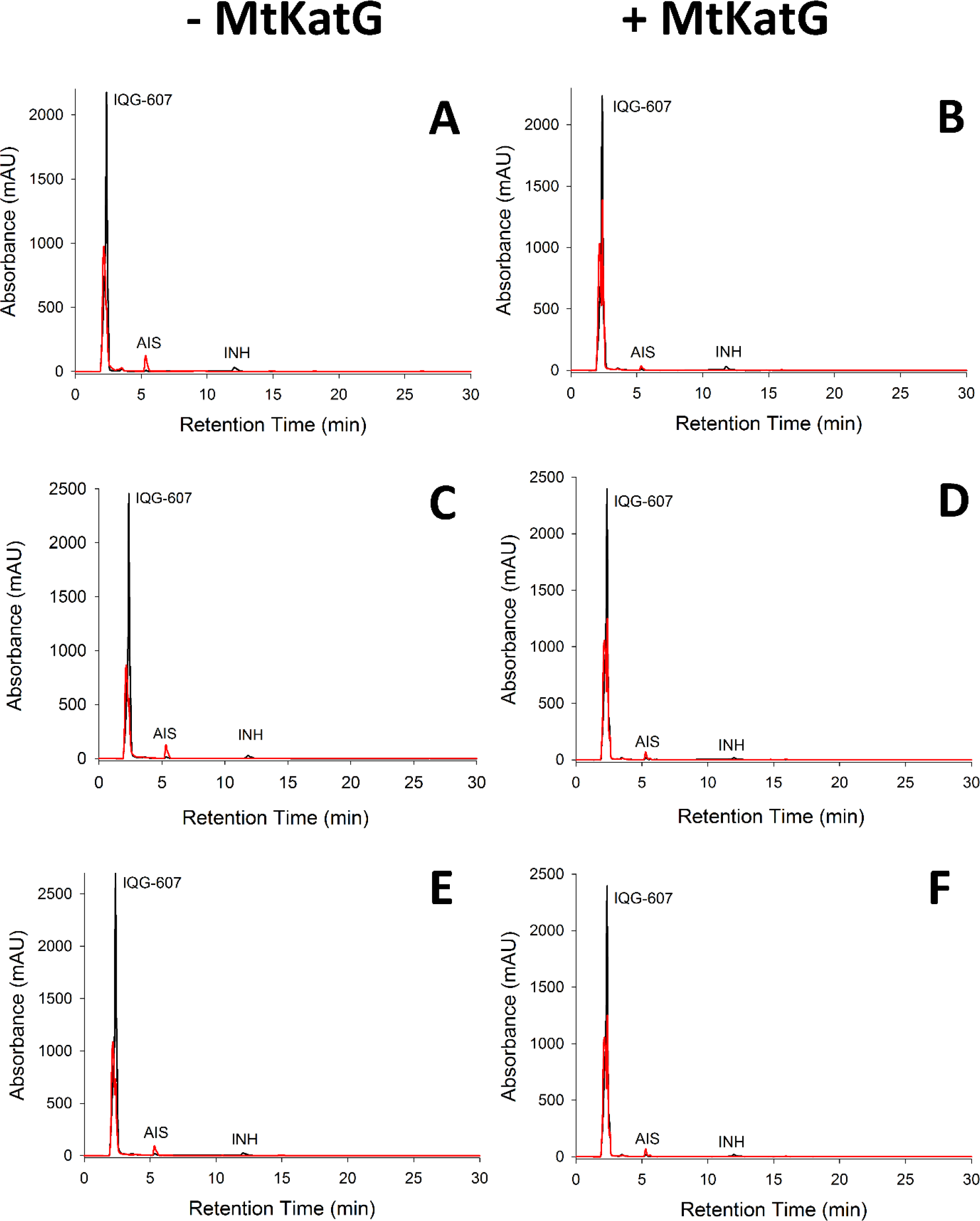
Representative chromatogram of IQG-607 with no enzyme in the reaction (-MtKatG) and addition of recombinant MtKatG (+ MtKatG). Oxidizing agents: air (a, b), H2O2 (C, D), and t-BuOOH (e, f). A major quantity of ISA is produced in the absence of MtKat

**Fig. 4.**
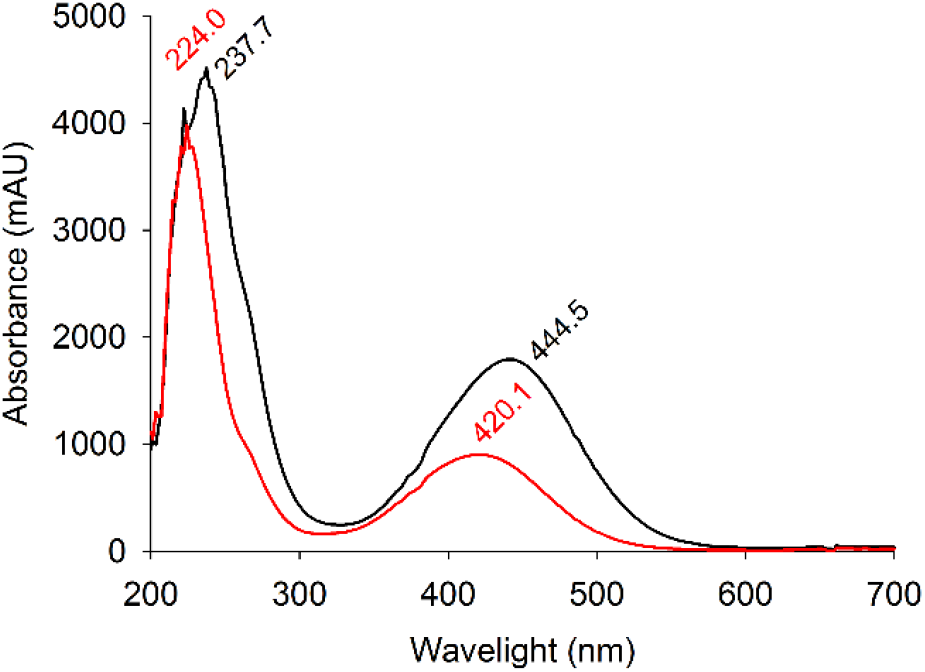
UV-spectra of IQG-607 over time. Time of incubation: t = 0 (black line), t = 100 min (red line).

**Table 3.**
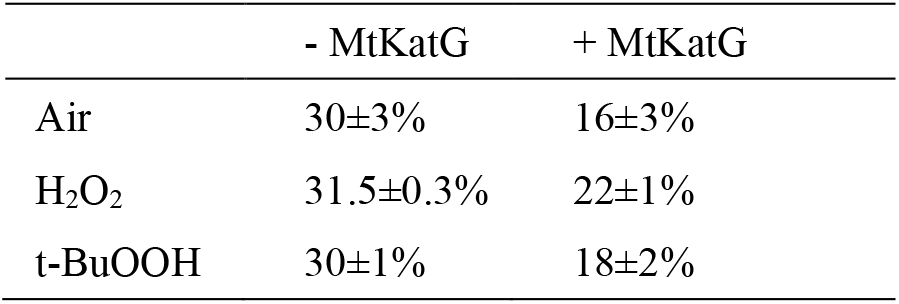
Percentage of IQG-607 degradation in the absence or presence of recombinant MtKatG.

|  | - MtKatG | + MtKatG |
| --- | --- | --- |
| Air | 30±3% | 16±3% |
| H<sub>2</sub>O<sub>2</sub> | 31.5±0.3% | 22±1% |
| t-BuOOH | 30±1% | 18±2% |

### INH and IQG-607 adduct formation with NAD^+^ using MtKatG

When using INH as a positive control, we observe that substrate consumption increased in the presence of oxidizing agents (Table 4). Not only ISA and INA are formed when INH reacts with MtKatG. In the presence of NAD^+^, the INH-NAD adduct is also formed, which is responsible for InhA inhibition inside Mtb cells, leading to cell death. We detect four INH-NAD adduct isomers (Fig. S4a-c). In the presence of t-BuOOH, MtKatG catalyzes the conversion of NAD^+^ to ADP-Ribose (Fig. S4c) and eventually to other products. However, INH-NAD adduct levels are unchanged in this condition when compared to the correspondent levels in hydrogen peroxide or air treatments. To confirm the identity of these adduct isomers, we synthesized the INH-NAD adduct using Mn(III) pyrophosphate (Fig. S4d), as previously reported [37,38].

**Table 4.** Percentage of consumption of INH in the absence or presence of recombinant MtKatG using NAD^+^ (adduct formation).

|  | - MtKatG | + MtKatG |
| --- | --- | --- |
| **Air** | ND* | 1.6±0.4% |
| **H<sub>2</sub>O<sub>2</sub>** | ND* | 2.2±0.8% |
| **t-BuOOH** | ND* | 5.8±0.9% |

\*ND: no degradation.

The HPLC profile of synthesized INH-NAD adduct isomers was then compared to the HPLC profiles obtained from adduct isomers enzymatically obtained using MtKatG in reactions containing INH and NAD^+^ as substrates and the absorption spectra of the INH-NAD adduct isomers from MtKatG-catalyzed reactions and synthesized adducts were very similar (Fig. S5). The identities of INH-NAD adduct obtained synthetically were confirmed by MS analysis (see Methods section 2.3). As expected, a parent ion of m/z 771.1535 corresponding to INH-NAD was detected and validated by detecting MS2 fragment ion products (Fig. S6). These results show that, apart from its catalase and peroxidase activities, the recombinant MtKatG can also produce the INH-NAD adduct in the presence of NAD^+^ and INH. In the presence of IQG-607, NAD^+^, and without MtKatG, there is no formation of INH-NAD (Fig. 5a, c, e). We detected a minor formation of INH-NAD adducts in reactions containing IQG-607, NAD^+^, and MtKatG, regardless of the oxidizing agent used (air-only, H_2_O_2_, or t-BuOOH) (Fig. 5b, d, f). Moreover, we observed again that MtKatG protects IQG-607 from decomposition (Table 5), as already observed in reactions without the addition of NAD^+^ (Table 3). Interestingly, in the presence of MtKatG and NAD^+^, t-BuOOH protects IQG-607 from decomposition (Fig. 5f and Table 5). It is important to notice that there is always residual INH (∼2%) from IQG-607 synthesis and we tried several methods to remove this residual INH without success. So, we supposed that the minor fractions of INH-NAD adduct isomers detected are derived from residual INH present in our reactions. We prepared control reactions spiked with different amounts of INH (25% and 50%) to confirm this hypothesis. Indeed, when compared to these control reactions, the amount of INH-NAD adducts obtained in reactions without spiking was proportional to the residual INH contained in those reactions (Fig. S7).

**Fig. 5.**
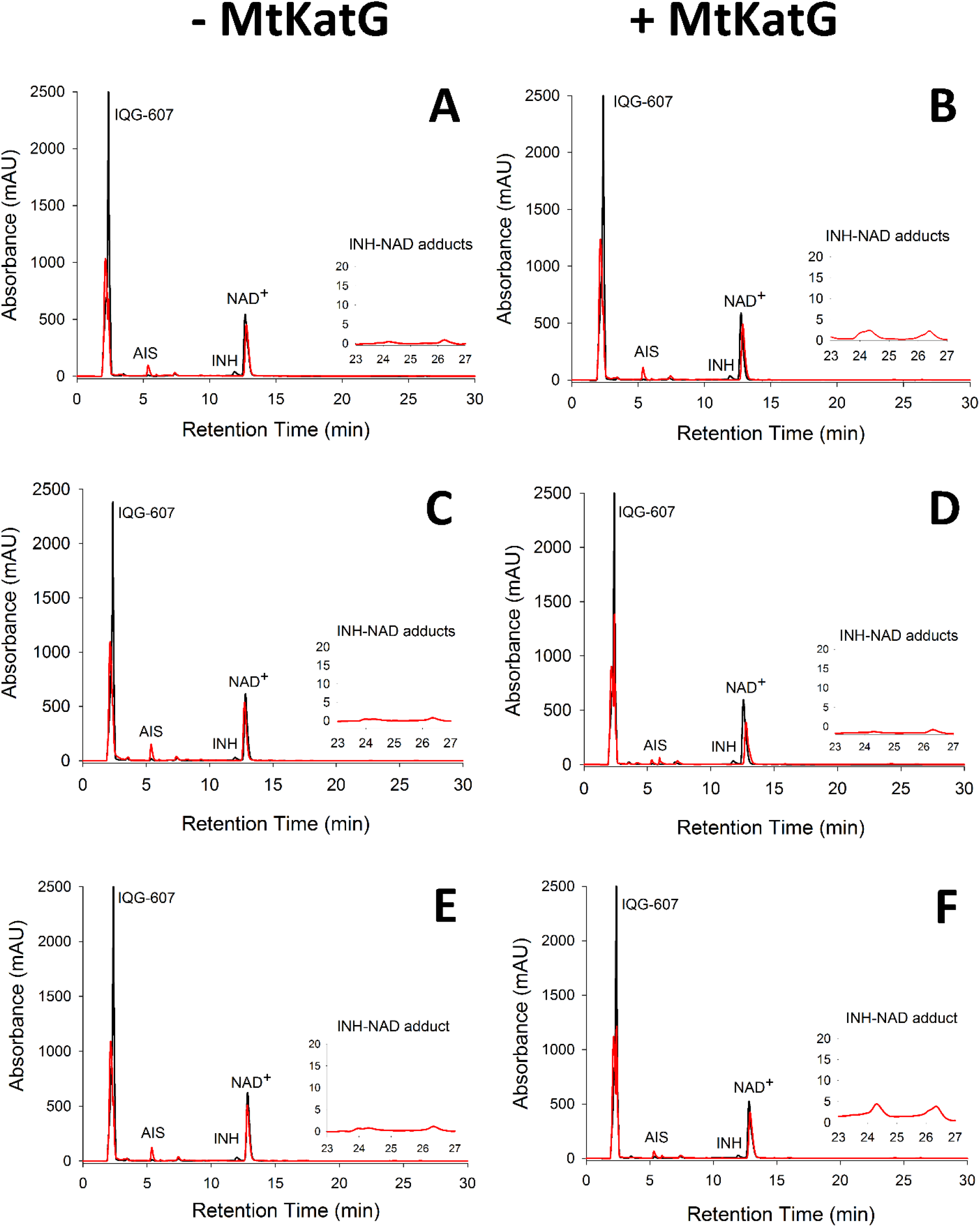
IQG-607 and NAD+ in absence (-MtKatG) or presence (+ MtKatG) of recombinant enzyme. Oxidizing agent: air (a, b), H_2_O_2_ (c, d), or t-BuOOH (e, f). The INH-NAD adducts were detected from 23 to 27 min (highlighted boxes). Time of incubation: t = 0 (black line), t = 100 min (red line).

**Table 5.** Percentage of degradation of IQG-607 in the absence and presence of recombinant MtKatG using NAD^+^.

|  | - MtKatG | + MtKatG |
| --- | --- | --- |
| **Air** | 29±1% | 27±1% |
| **H<sub>2</sub>O<sub>2</sub>** | 33±1% | 21±5% |
| **t-BuOOH** | 25±3% | 11±4% |

### Puzzle of IQG-607’s Mechanism of Action

Metal-based drugs are attractive therapeutic agents that are increasingly being used in a broad range of medical conditions, both for therapy and diagnosis [16]. However, the detailed molecular mechanisms of action of these compounds are still elusive. This knowledge is instrumental in the tailoring of metallodrugs to specific therapeutic purposes.

The metal-based compound IQG-607 is an INH derivative containing a pentacyanoferrate(II) attached to INH. It was developed to bypass the requirement for KatG activation during the intracellular production of the active INH-NAD adduct from the INH prodrug. Such compounds could be used eventually to treat TB infections caused by INH-resistant isolates whose resistance mechanism relies on mutations on the *katG* gene, a significant cause of INH resistance in clinics. We have previously shown that inside mycobacterial cells, the antimicrobial activity of the IQG-607 requires an active KatG enzyme [27]. This finding does not support the metal-based autoactivation mechanism originally envisaged for this molecule [20, 21]. In the title of our previous review of the compound IQG-607, we asked whether this molecule is a metallodrug or a metallopro-drug [48]. Based on the available evidence at the time, we ended the study positing that IQG-607 would act as a metallopro-drug, requiring KatG enzyme to form an active drug inside mycobacterial cells. However, the present work shows that there is no formation of in vitro INH adducts. After assaying a recombinant form of MtKatG in the presence of NAD^+^ and different oxidizing agents, we found no evidence that IQG-607 could serve as a substrate for adduct formation under all conditions tested. The results obtained in this work may support an alternative hypothesis to explain why IQG-607 requires KatG activation to exert its antimicrobial effects. This compound might behave as a pre-prodrug, releasing the INH moiety inside mycobacterial cells, requiring KatG activation to form the active INH-NAD adduct. In this way, efforts need to be made to support this hypothesis. So, the current work adds another piece to the puzzle of IQG-607’s mechanism of action. Using a combination of genetic [27] and biochemical approaches [this study], we investigated the mechanisms underpinning the antimicrobial activities of the metal-based compound IQG-607. A similar approach could be employed to study other metallodrugs of therapeutic value as antimicrobial agents.

## Abbreviations

Mtb: *Mycobacterium tuberculosis*
InhA: Enoyl-ACP-reductase KatG-Catalase-peroxidase
TB: Tuberculosis
t-BuOOH: Tert-butyl hydroperoxide
RIF: Rifampicin
RR: Rifampicin resistance
INH: Isoniazid
EMB: Ethambutol
PZA: Pyrazinamide
MDR: Multidrug-resistant
XDR: Extensively drug-resistant
NAD^+^: Nicotinamide adenine dinucleotide
ROS: Reactive oxygen species
WT: Wild type
LMW: Low molecular weight
HMW: High molecular weight
ABTS: 2,2’-azino-bis(3-ethylbenzothiazoline-6-sulfonic acid
ISA: Isonicotinic acid
INA: Isonicotinamide

## Declarations

### Availability of data and material

Chromatography data are available upon request.

### Code availability

Not applicable.

### Authors’ Contributions

LQB performed experiments, analyzed data, and wrote the manuscript. CDR performed experiments (MtKatG purification and enzyme activity assays). ASD developed methods and performed experiments (development of HPLC methods and oxidation assays) RSR performed experiments (synthesis of IQG-607, spiking experiments) and wrote the manuscript. PFD performed experiments (LC-MS/MS analysis). KP performed experiments (synthesis of IQG-607, INH-NAD adduct). JESN performed experiments (gel filtration experiments). RJE performed experiments (MtKatG expression). PM designed experiments and revised the manuscript. LAB designed experiments and revised the manuscript. CVB designed experiments, analyzed data, and wrote the manuscript.

