## Supplementary materials for "The isoniazid analog IQG-607 is not a direct substrate for the *Mycobacterium tuberculosis* catalase-peroxidase KatG"

a. Instituto Nacional de Ciência e Tecnologia em Tuberculose (INCT-TB), Centro de Pesquisas em Biologia Molecular e Funcional (CPBMF), Pontifícia Universidade Católica do Rio Grande do Sul (PUCRS), 92A TECNOPUC, 4592 Av. Ipiranga 6681, 90616-900, Porto Alegre, Brazil.

b. Programa de Pós-Graduação em Biologia Celular e Molecular, PUCRS, Porto Alegre, Brazil.

c. Programa de Pós-Graduação em Medicina e Ciências da Saúde, PUCRS, Porto Alegre, Brazil


*E. coli* BL21 (DE3)pLysS cells were transformed with pET23a(+)::*katG* were grown on Lysogenic Broth (LB) agar plates containing 50 mg mL$^{-1}$ ampicillin and 34 mg mL$^{-1}$ chloramphenicol. A single colony was cultivated overnight in 50 mL of LB at 37 ºC and 180 rpm. After that, 10 mL of the culture were inoculated into 500 mL of LB medium with the same concentrations of antibiotics and grown at 37 ºC and 180 rpm until an $OD_{600}$ of 0.5. To induce MtKatG expression, IPTG was added to a final concentration of 1 mM, together with 30 mg L$^{-1}$ of Hemin, to ensure stoichiometric incorporation of the heme protestic group. Cells were grown for 6 hours and harvested by centrifugation at 11000 *g* for 30 min at 4 ºC and stored for several months at - 20 ºC. To obtain a purified MtKatG, 2 g of frozen cells were resuspended in 20 mL of 50 mM Tris HCl pH 7.6 (buffer A) containing 0.2 mg mL$^{-1}$ of lysozyme and stirred for 30 min. Cells were disrupted by sonication (10 pulses of 10 s each at 60% amplitude) and centrifuged at 38900 *g* for 30 min. The supernatant was incubated with 1% (v/v) of streptomycin sulfate for nucleic acid removal and gently stirred for 30 min. The solution was centrifuged at 38900 *g* for 30 min. The supernatant was dialyzed two times against 2 L of buffer A using a dialysis tubing with a molecular weight exclusion limit of 12000 – 14000 Da. After the dialysis, the solution was centrifuged at 38900 *g* for 30 min and the supernatant was loaded on a Q Sepharose Fast Flow (GE Healthcare) pre-equilibrated with buffer A. The column was washed with 5 column volumes (CV) of the same buffer, and adsorbed proteins were eluted with a linear gradient (0 – 100%) of 15 CV of 50 mM Tris HCl pH 7.6 containing 1 M NaCl (buffer B) at 3 mL min$^{-1}$ flow rate. The recombinant MtKatG protein eluted between 250 and 330 mM of NaCl concentration. Fractions containing the target protein were pooled and ammonium sulfate was added to a final concentration of 1 M, clarified by centrifugation at 38900 *g* for 30 min, and the resulting supernatant was loaded on a HiLoad 16/10 Phenyl Sepharose High Performance (GE Healthcare) pre-equilibrated with 50 mM Tris HCl pH 7.6 containing 1 M $(NH_4)_2SO_4$ (buffer C). This hydrophobic column was washed with 5 CVs of buffer C and the adsorbed material eluted with 20 CVs of a linear gradient (0 – 100%) of buffer A at 1 mL min$^{-1}$ flow rate. The protein eluted between 425 mM and 342 mM of buffer C. The fractions containing the KatG protein were pooled, concentrated down to 5 mL using an Amicon ultrafiltration cell (molecular weight cutoff of 30000 Da), and loaded on a size exclusion column HiLoad Superdex 200 (GE Healthcare), which was previously equilibrated with 100 mM potassium phosphate pH 7.5 (buffer D). Proteins were isocratically eluted with 1 CV of buffer D at a flow rate of 0.5 mL min$^{-1}$. Protein concentration was determined by the method of BCA using bovine serum albumin as standard (Thermo Scientific Pierce™ BCA protein Assay Kit).

```
001 MPEQHPPITE TTTGAASNGC PVVGHMKYPV EGGGNQDWWP NRLNLKVLHQ NPAVADPMGA 060
061 AFDYAAEVAT IDVDALTRDI EEVMTTSQPW WPADYGHYGP LFIRMAWHAA GTYRIHDGRG 120
121 GAGGGMQRFA PLNSWPDNAS LDKARRLLWP VKKKYGKKLS WADLIVFAGN CALESMGFKT 180
181 FGFGFGRVDQ WEPDEVYWGK EATWLGDERY SGKRDLENPL AAVQMGLIYV NPEGPNGNPD 240
241 PMAAAVDIRE TFRRMAMNDV ETAALIVGGH TFGKTHGAGP ADLVGPEPEA APLEQMGLGW 300
301 KSSYGTGTGK DAITSGIEVV WTNTPTKWDN SFLEILYGYE WELTKSPAGA WQYTAKDGAG 360
361 AGTIPDPFGG PGRSPTMLAT DLSLRVDPIY ERITRRWLEH PEELADEFAK AWYKLIHRDM 420
421 GPVARYLGPL VPKQTLLWQD PVPAVSHDLV GEAEIASLKS QIRASGLTVS QLVSTAWAAA 480
481 SSFRGSDKRG GANGGRIRLQ PQVGWEVNDP DGDLRKVIRT LEEIQESFNS AAPGNIKVSF 540
541 ADLVVLGGCA AIEKAAKAAG HNITVPFTPG RTDASQEQTD VESFAVLEPK ADGFRNYLGK 600
601 GNPLPAEYML LDKANLLTLS APEMTVLVGG LRVLGANYKR LPLGVFTEAS ESLTNDFFVN 660
661 LLDMGITWEP SPADDGTYQG KDGSGKVKWT GSRVDLVFGS NSELRALVEV YGADDAQPKF 720
721 VQDFVAAWDK VMNLDRFDVR
```

**Figure S1.** Peptide mapping of recombinant MtKatG (GenBank: AVV29661.1). The primary structure is represented and peptides in blue were identified by LC-MS/MS. A 91% coverage were obtained.

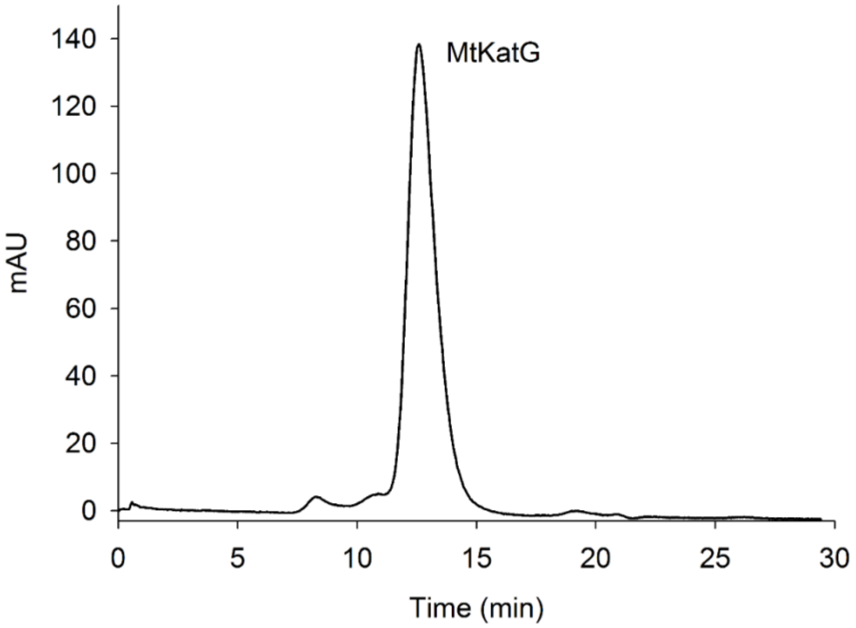


**Figure S2.** Determination of the oligomeric state of recombinant MtKatG by gel filtration using a HighLoad 10/30 Superdex-200 column. Protein elution profile at 215 nm.

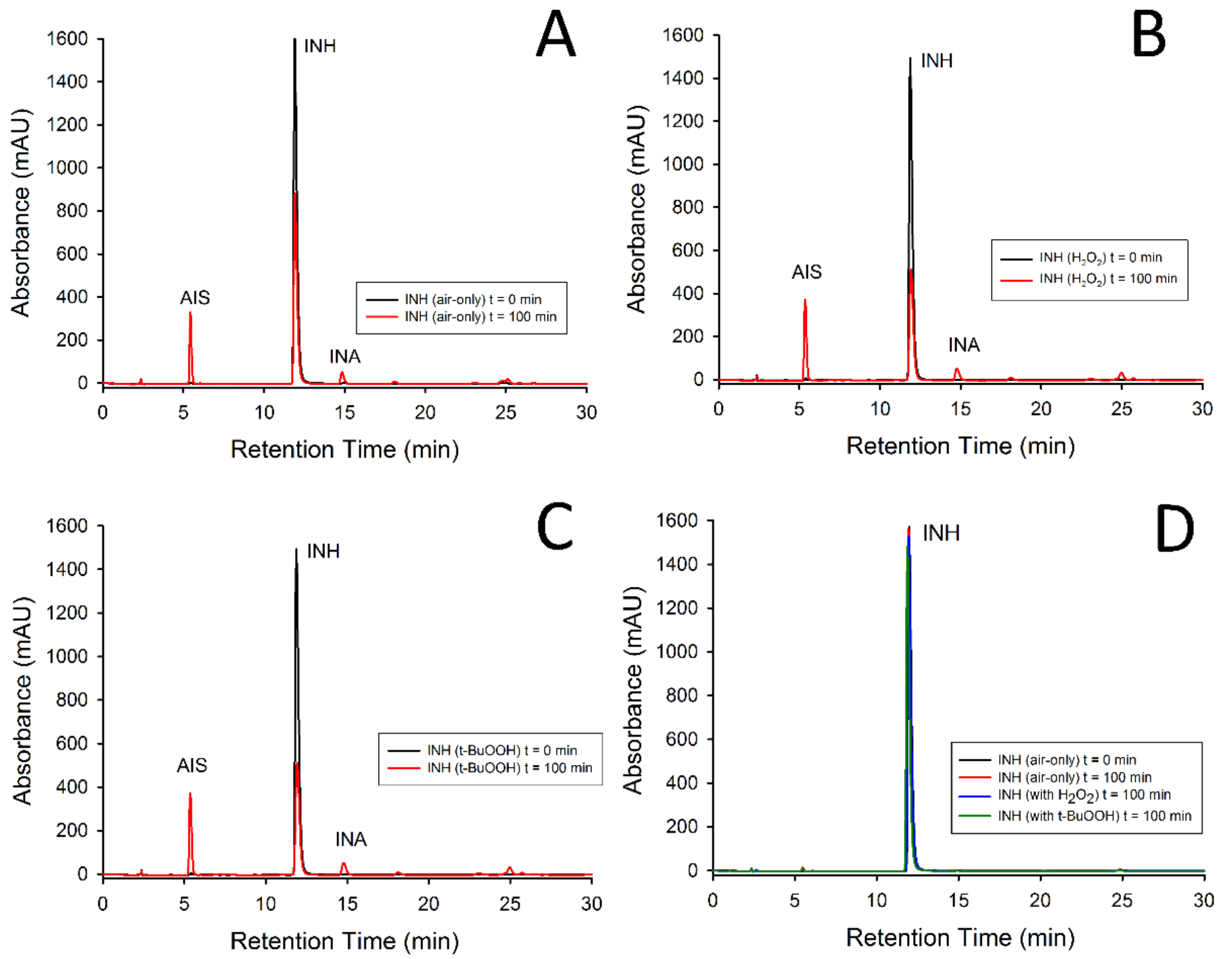


**Figure S3.** INH in presence of recombinant MtKatG. Background air were used as oxidant agent (a), $H_2O_2$ (b) and t-BuOOH (c) as peroxidative agents. The black line represents t = 0 min of incubation and red line represents t = 100 min of incubation. INH is also tested in absence of recombinant MtKatG and showed no degradation into products, even in addition of peroxides (d).

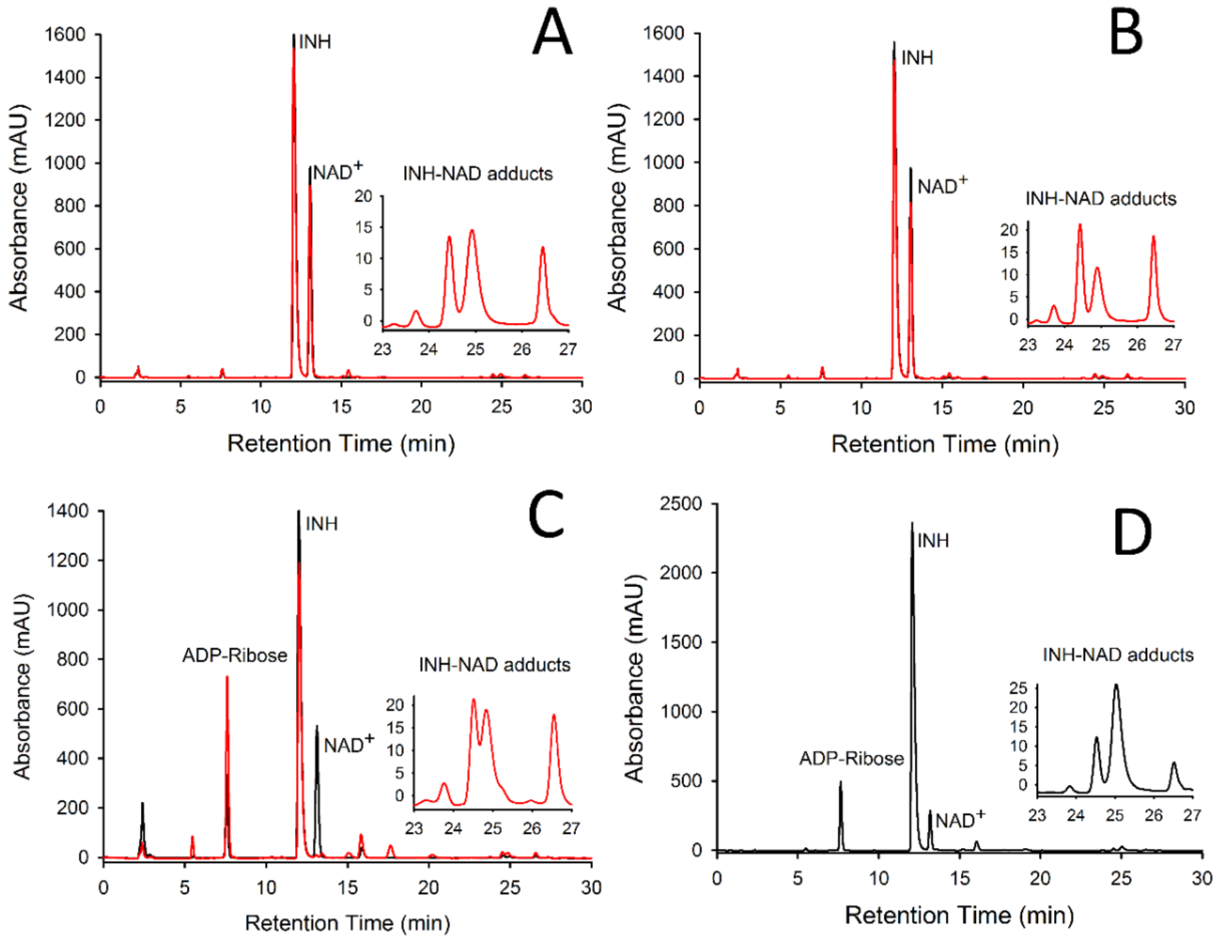


**Figure S4.** INH and NAD$^+$ in presence of recombinant MtKatG. The INH-NAD adducts were detected between 23 and 27 min. The chromatograms represent when air-only were used (a), $H_2O_2$ (b) and t-BuOOH (c) as peroxidative agents. The black line represents t = 0 min of incubation and red line represents t = 100 min of incubation. Chemical synthesis of INH-NAD using Mn (III) pyrophosphate (d).

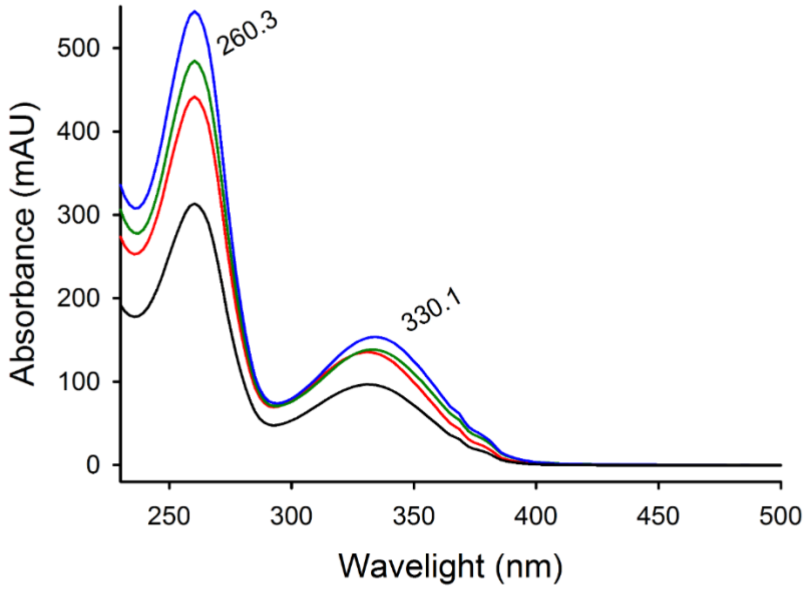


**Figure S5.** Absorption spectroscopy of INH-NAD analysis. The four isomers identified in HPLC-UV showed the same profile (260/330).

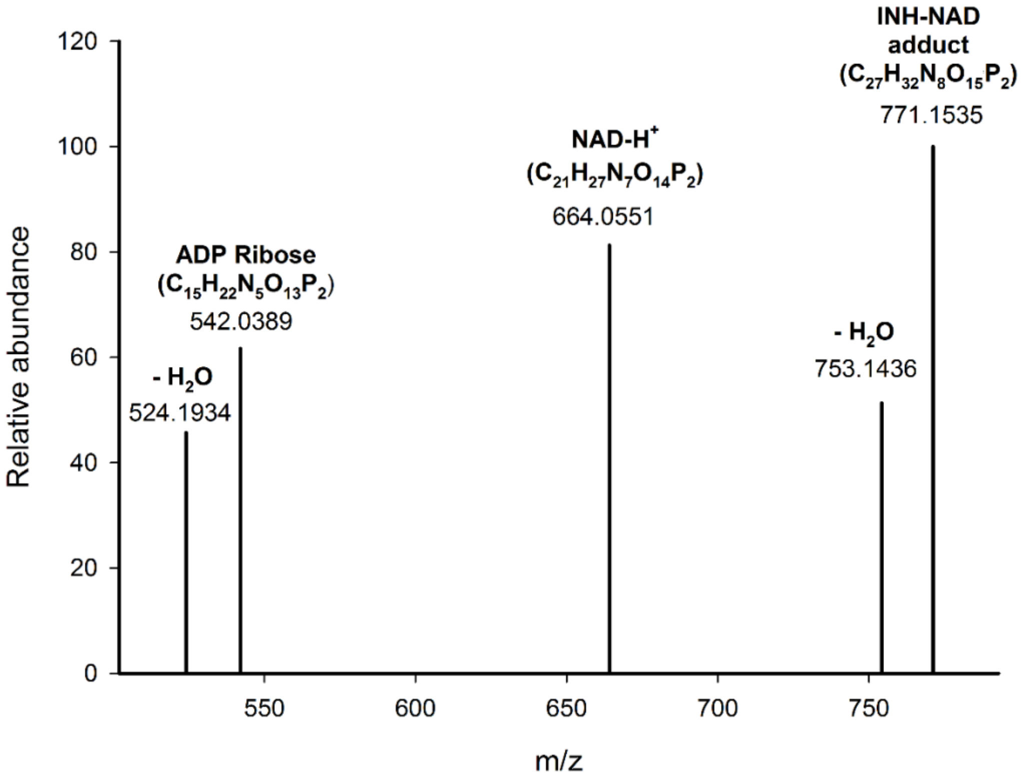


**Figure S6.** Identification of INH-NAD by LC-MS/MS. The spectra of m/z = 771.1535 correspond to INH-NAD adduct, and a water loss also can be observed (m/z = 753.1436). NAD-$H^+$ spectra of m/z = 664.0551 and ADP Ribose with a water loss can also been observed (m/z of 542.0389 and 524.1934, respectively).

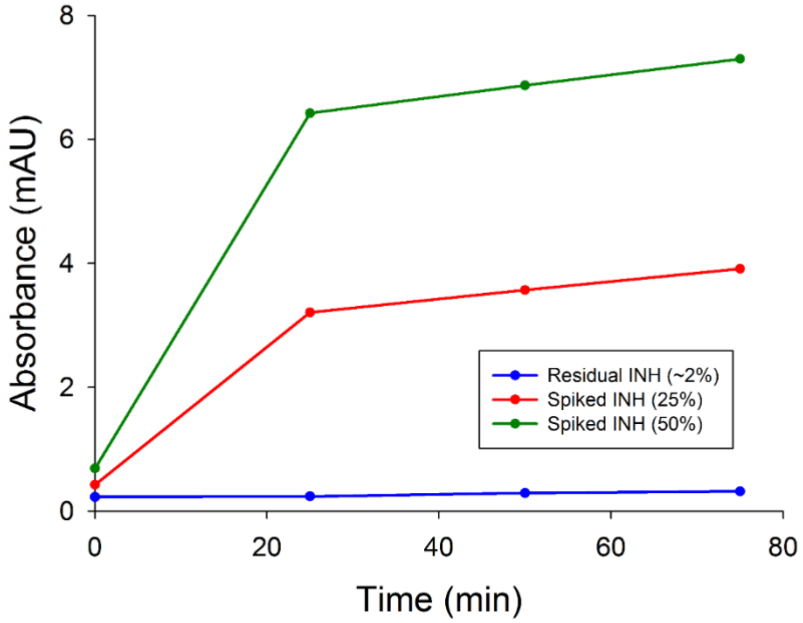


**Figure S7.** Formation of INH-NAD when INH were spiked in IQG-607. The increase in absorbance of INH-NAD adduct (mAU) over time were observed when INH were spiked in IQG-607 reaction, demonstrating that INH-NAD formation comes from residual INH.
